# Function and evolution of aquaporin IsAQP1 in the Lyme disease vector *Ixodes scapularis*

**DOI:** 10.1101/2022.07.26.501577

**Authors:** Hitoshi Tsujimoto, Hillery C. Metz, Alexis A. Smith, Joyce M. Sakamoto, Utpal Pal, Jason L. Rasgon

**Affiliations:** Department of Entomology, Center for Infectious Disease Dynamics and the Huck Institutes of the Life Sciences, Pennsylvania State University, University Park, PA, 16832, USA; Department of Entomology, Texas A&M University, College Station, TX, 77843, USA; Department of Veterinary Medicine, University of Maryland, College Park, MD, 20742, USA

**Author notes:** Correspondence: Jason L. Rasgon, Pennsylvania State University, University Park, PA, 16832.

**Keywords:** *Ixodes*, ticks, aquaporins, Lyme disease, osmoregulation

## Abstract

Ticks are important vectors of pathogenic viruses, bacteria, and protozoans to humans, wildlife, and domestic animals. Due to their life cycle, ticks face significant challenges related to water homeostasis. When blood-feeding they must excrete water and ions, but when off-host (for stretches lasting several months) they must conserve water to avoid desiccation. Aquaporins (AQPs), a family of membrane-bound water channels, are key players in osmoregulation in many animals but remain poorly characterized in ticks. Here, we bioinformatically identified AQP-like genes from the deer tick *Ixodes scapularis* and used phylogenetic approaches to map the evolution of the aquaporin gene family in arthropods. We find that most arachnid AQP-like sequences (including those of *I. scapularis*) form a monophyletic group clustered within aquaglycerolporins (GLPs) from bacteria to vertebrates. This gene family is absent from insects, revealing divergent evolutionary paths for AQPs in different hematophagous arthropods. We next sequenced the full-length cDNA of *I. scapularis* aquaporin 1 (IsAQP1) and expressed it heterologously in *Xenopus* oocytes to functionally characterize its permeability to water and solutes. We additionally examined IsAQP1 expression across different life stages and adult female tissues. We found IsAQP1 to be an efficient water channel with salivary gland-specific expression, implying its function may be closely tied to osmoregulation during blood-feeding. Its functional properties were unique: unlike most GLPs, IsAQP1 has low glycerol permeability, and unlike most AQPs, it is insensitive to mercury. Together, our results suggest IsAQP1 plays an important role in tick water balance during blood feeding and may hold promise as a target for novel vector control efforts.

**Graphical TOC/Abstract:** 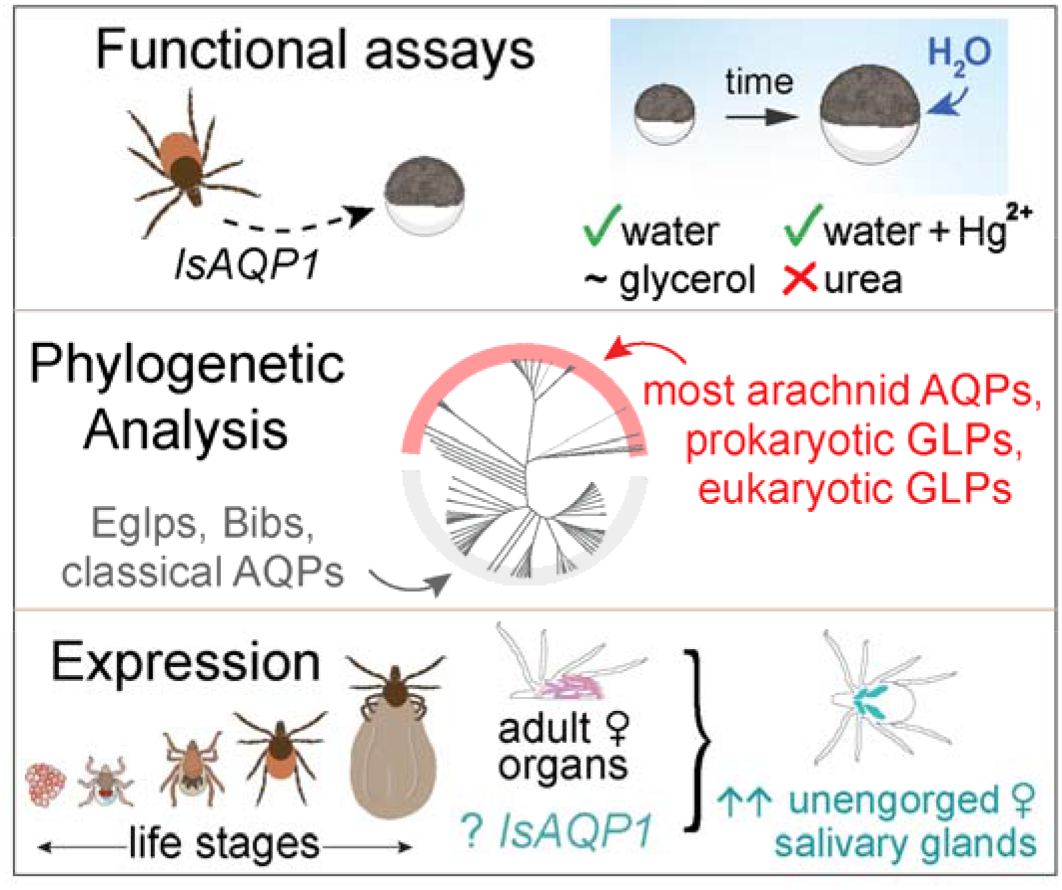

*eTOC:* IsAQP1, an aquaporin found in the black legged tick (*Ixodes scapularis*), is a functional water channel with low permeability to glycerol. In a phylogenetic analysis of 176 sequences from 48 arachnid species, most tick and other arachnid AQP-like genes formed a monophyletic clade nested within aquaglyceroporins (GLPs). *IsAQP1* shows salivary-gland specific expression in adult females—with highest expression in unfed females—suggesting a role in water balance during blood feeding.

## Introduction

Ticks are obligate blood-feeding arachnids that are not only pests of livestock and humans, but also vectors of numerous pathogens including viruses, bacteria, protozoans, and nematodes. The deer tick, *Ixodes scapularis*, is the most important vector of pathogens in North America and is a major vector of *Borrelia burgdorferi*, the Lyme disease agent. The increasing burden of tick-borne diseases—on people and animals alike—necessitates new methods for controlling this pernicious vector (Dantas-Torres *et al*., 2012; Boulanger *et al*., 2019). However, an obstacle to developing new tools is a lack of molecular detail in our current understanding of tick biology.

Ticks are long-lived arthropods (living up to several years) with a multistage life history that creates significant and opposing water balance challenges. On one hand, while attached to hosts, ticks must excrete excess water and ions from the host blood they consume—a major physiological challenge because a blood meal may be 100 times their unfed body weight (Kaufman and Phillips, 1973). This excess water is removed via salivary secretion back into the host (Bowman and Sauer, 2004). On the other hand, ticks face desiccation risk during the extended periods they spend off-host (Benoit and Denlinger, 2010), during which water can only be obtained by absorbing it from the air into salivary tissues (Needham and Teele, 1991). ‘Three-host ticks’ including *I. scapularis* require a host blood meal during each of three distinct life stages, with extended periods spent living off-host without feeding between these meals (Boulanger *et al*., 2019). The water homeostasis needs of *I. scapularis* therefore fluctuate dramatically across life stages, and these fluctuating needs may be reflected in stage- or tissue-specific expression of osmoregulatory molecules.

Aquaporins (AQPs) are a family of tetrameric transmembrane water channels found in the genomes of most organisms, from bacteria to plants and animals (Abascal *et al*., 2014). They facilitate the movement of water across cells and membranes and have traditionally been classified as either water-specific (‘classical aquaporins’), or as additionally permeable to small molecules such as glycerol or urea (the aquaglyceroporins, or GLPs; Abascal *et al*., 2014). Generally, but not always, these traditional functional categories comport with AQP gene trees—i.e., phylogenetic analysis results in distinct clades of AQPs and GLPs (e.g., Finn *et al*., 2015; Tsujimoto *et al*., 2017). The permeabilities and other properties of each channel are determined by a handful of residues within the channel pore, and changes to a few amino acids can alter channel permeability (Fu *et al*., 2000; Sui *et al*., 2001; Beitz *et al*., 2006, Campbell *et al*. 2008). Indeed, mismatches between measured channel function and traditional categories (i.e., classical AQP or GLP) have been described. For example, insect ‘Eglps’ have GLP-like permeability to solutes yet they evolved from classical water-specific AQPs (Finn *et al*., 2015), and Big brain (Bib) gene products appear to have lost permeability to water and other substrates but gained a new role in embryonic development (Rao *et al*., 1990; Doherty *et al*., 1997).

As water channels, aquaporins mediate essential homeostatic processes including water balance (e.g., Liu *et al*., 2011; Drake *et al*., 2010, 2015). Glycerol conductance (e.g., by GLPs) can also be important for survival (Philip *et al*., 2008, 2011), as some arthropods accumulate glycerol and other polyols to survive cold or desiccating conditions (e.g., Duman, 2001; Yoder *et al*., 2006). Knockdown of AQPs has been shown to negatively impact hematophagous insects. In one case, reduction of an AQP and an Eglp decreased the ability of the bed bug, *Cimex lectularius*, to excrete waste following a blood meal (Tsujimoto *et al*., 2017). Other studies functionally verified a role for AQPs in modulating desiccation status in the mosquitoes *Anopheles gambiae* and *Aedes aegypti* (Drake *et al*., 2010; Liu *et al*., 2011), and in osmoregulation in the ticks *Ixodes ricinus* and *Rhipicephalus microplus* (Campbell *et al*. 2010; Hussein *et al*. 2015). Outside of insects, less is known about arthropod aquaporins (but see Stavang *et al*., 2015, Benoit *et al*. 2014b, Campbell *et al*. 2008). Of the >700 hard tick and nearly 200 soft tick species (Boulanger *et al*., 2019), an aquaporin channel has thus far been functionally characterized in only one: *Rhipicephalus sanguineus* (Ball *et al*., 2009). We therefore studied putative AQP-like genes in *I. scapularis*, examined the pattern of AQP evolution in arthropods more broadly, and characterized both the functional properties and tissue- and stage-specific expression patterns of the *I. scapularis* aquaporin IsAQP1.

## Results

### Identification of *I. scapularis* aquaporins and characterization of *IsAQP1*

We identified 18 putative aquaporin-like sequences for *Ixodes scapularis* including ISCW003957 (IsAQP1; Table S1). We next acquired the full-length cDNA sequence for one of these genes, *Ixodes scapularis* aquaporin-1 (IsAQP1) using RACE. The *IsAQP1* mRNA consists of 1,481 nucleotides (excluding the poly-A tail) with a 885-nt coding sequence encoding a 294-aa polypeptide. When mapped to the genome, it spans 62,141 nucleotides on the Scaffold:IscaW1:DS692803 positive strand and comprises 7 exons and 6 introns (Figure 1). Our analysis adds two exons (one to the 5’ end and one to the 3’ end) to a previous genome annotation by VectorBase (geneset: IscaW1.5). The start and stop codons are located in these new exons, which is now reflected in the geneset IscaW1.6. The sequence information was submitted to GenBank with an accession number KT988052. The polypeptide contains two NPA motifs—a conserved feature of AQPs (Murata *et al*., 2000; Benga 2012) —complete with RD residues after the second NPA motif uniquely conserved in aquaglyceroporins (GLPs) (Fu *et al*., 2000; Sui *et al*., 2001).

**Figure 1.**
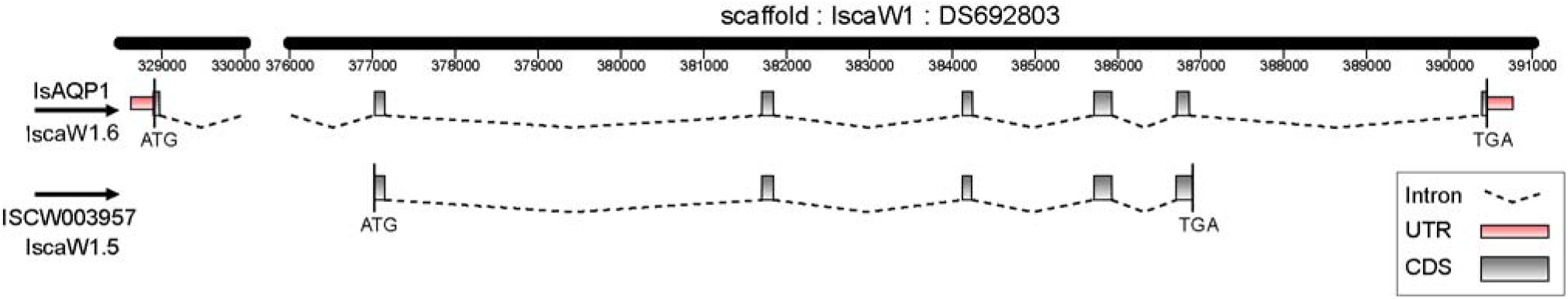
ISCW003957 (*IsAQP1*) gene structure in the *I. scapularis* genome. Top: the structure obtained by this study; bottom: the structure by the previous annotation (IscaW1.5). In both, start (ATG) and stop (TGA) codons are indicated.

### Phylogenetic analysis of arthropod aquaporins reveals distinct evolution in arachnids

We examined the relationship of the newly identified *I. scapularis* AQPs within the broader gene family using maximum likelihood phylogenetic analysis. We constructed a gene tree for AQP and AQP-like genes from diverse arthropods including hexapods, crustaceans, and chelicerates that includes 12 orders and 48 species (Figure 2). We additionally included 17 aquaporin sequences from outgroup taxa *E. coli, M. tardigradum*, and *H. sapiens*. The data set included 176 total AQP and AQP-like sequences. Most (68/94, 72%) arachnid AQPs (represented by *Aranae, Ixodida, Mesostigmata, Scorpiones*, and *Trombidiformes*) formed a clade nested within the classical aquaglyceroporins (GLPs). This pattern was even more pronounced among tick sequences: only six of 57 tick AQPs (10.5%) did not cluster within the arachnid GLP clade, five of which appeared to be Bib brain (Bib) genes. As expected, insect and crustacean (i.e., pancrustacean, Giribet and Edgecombe, 2019) AQPs and insect-specific Eglps both clustered with classical water-specific AQPs (Figure 2).

**Figure 2.**
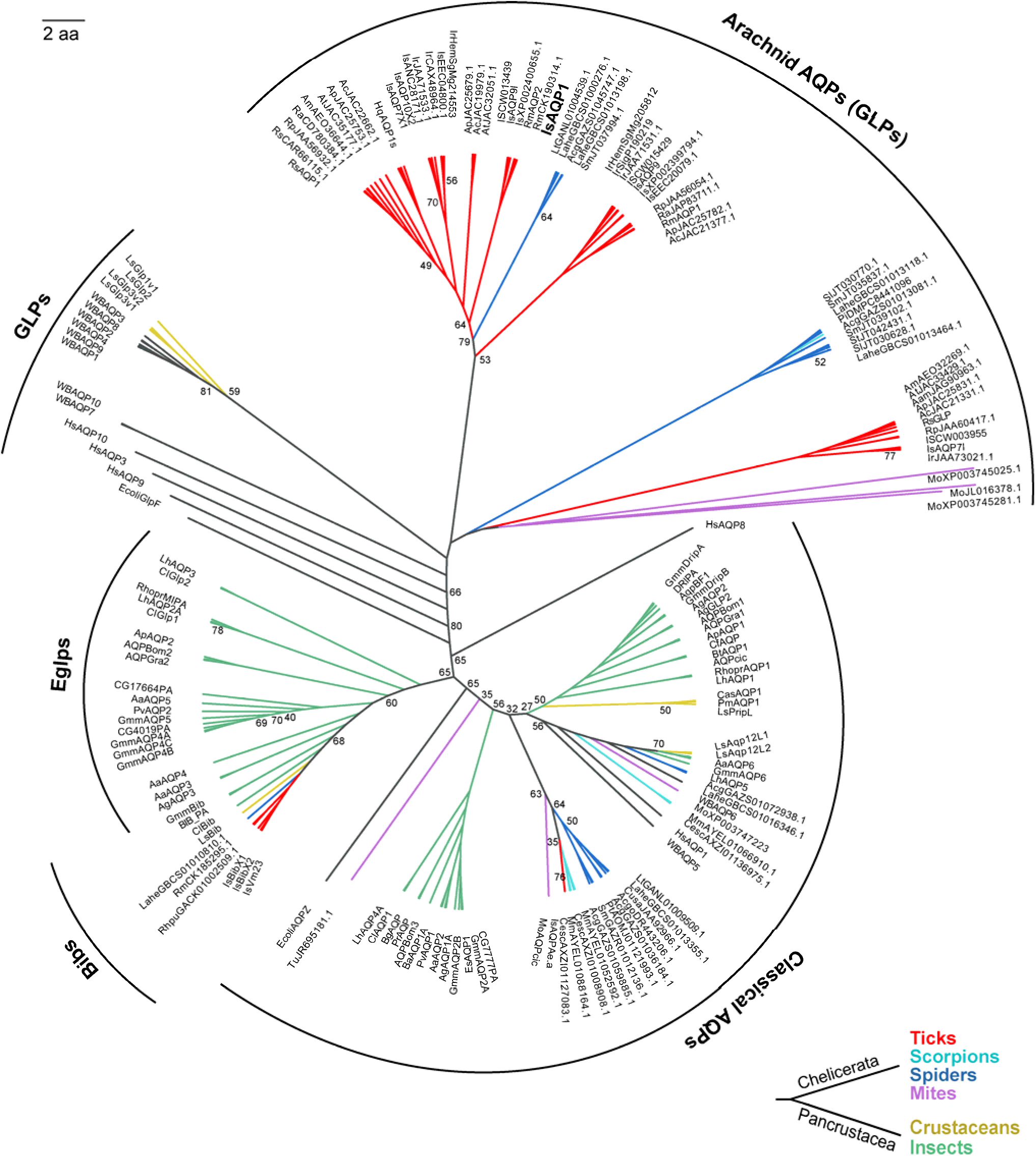
Phylogenetic tree of selected aquaporins (AQPs) and aquaglyceroporins (GLPs) from arthropods and other organisms. Bootstrap values (1000 replicates) are indicated at each node; unlabeled nodes have bootstrap values ≥85. Sequence IDs are as in Table S1. The scale bar represents amino acid substitutions.

### IsAQP1 is a functional water channel with limited glycerol permeability and no urea permeability

We expressed IsAQP1 heterologously in *Xenopus laevis* oocytes to assess its water and solute permeability. IsAQP1 channels exhibited significantly greater water permeability in comparison to vehicle (nuclease-free water) injected controls (Fig. 3A, one-way ANOVA with post hoc Tukey test, *P*<0.05). Indeed, IsAQP1 moved water more efficiently (1.47x) than the positive control classical AQP channel AgAQPB1 (Fig. 3A, one-way ANOVA with post hoc Tukey test, *P*<0.05). However, unlike many characterized AQPs (e.g., Hirano *et al*. 2010, Campbell *et al*. 2008), it was not significantly inhibited by Hg^2+^ (Fig. 3A, *P*>0.05). IsAQP1 also had low glycerol permeability (0.28x that of the positive control; Fig. 3B, one-way ANOVA with post hoc Tukey test, *P*<0.05). It was not permeable to urea (Fig. 3C; *P*>0.05).

**Figure 3.**
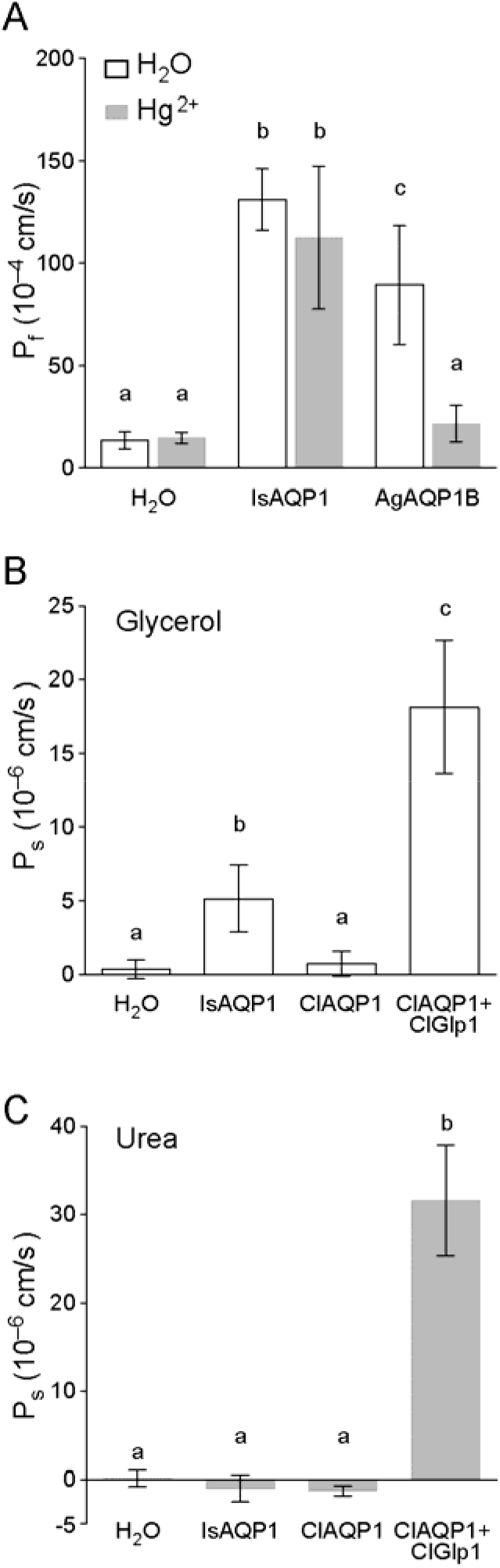
Water, glycerol and urea permeability of IsAQP1 (ISCW003957) alongside selected insect channels (positive controls). (A) Water permeability of *Xenopus* oocytes expressing IsAQP1 in the presence or absence of conventional inhibitor Hg^2+^. (B) Glycerol permeability. (C) Urea permeability. H_2_O: water-injected negative control; AgAQP1: *Anopheles gambiae* AQP1-expressing oocyte. ClAQP1: *Cimex lectularius* AQP1 expressing oocyte. ClAQP1+ClGlp1: *Cimex lectularius* AQP1 and Glp1 co-expressing oocyte. Letters indicate statistical grouping according to one-way ANOVA with *post hoc* Tukey tests.

### *IsAQP1* shows tissue and life-stage specific expression differences

We examined the expression pattern of IsAQP1 using quantitative PCR. We first measured transcript expression across life stages including egg, larva, nymph, and adults of both sexes. Eggs and nymphs showed minimal IsAQP1 expression, while expression at other stages (larva, adults) was higher on average, but highly variable across biological replicates (Fig. 4A). Expression in male and female adults was also characterized by high variation across replicates, but the sexes did not appear to differ (Fig. 4A). Due to high variability between replicates, we did not statistically compare IsAQP1 expression in these stages or sexes.

**Figure 4.**
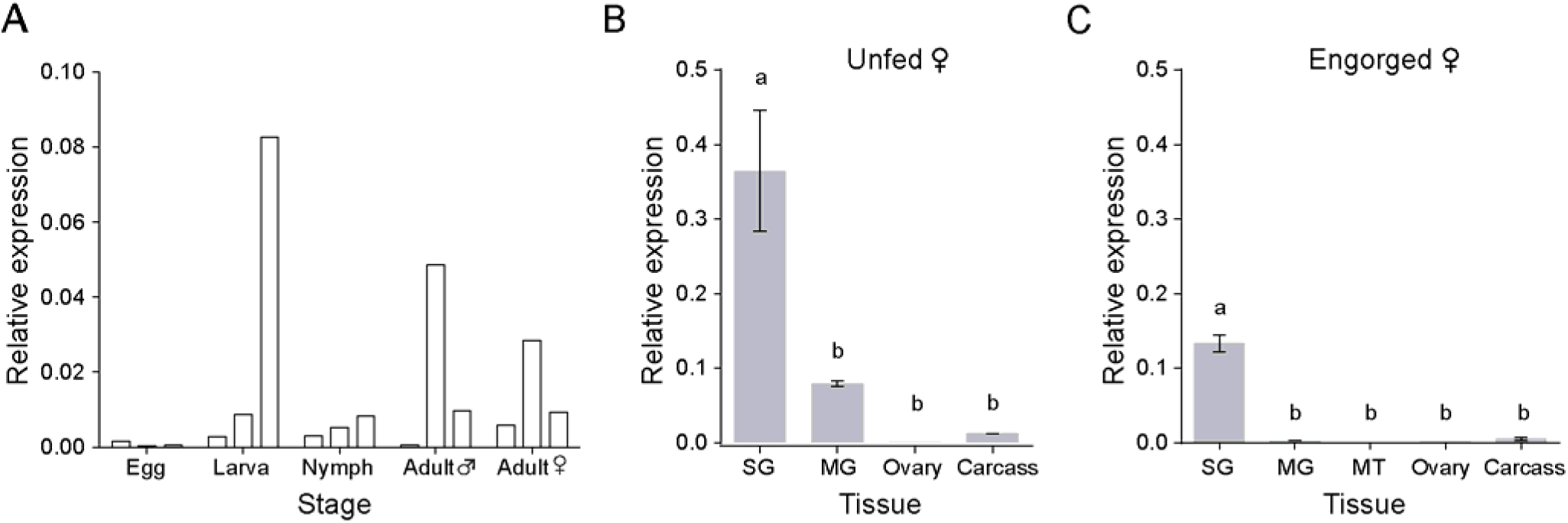
Quantification of *IsAQP1* mRNA expression by qRT-PCR. (A) *IsAQP1* expression across several developmental stages. Each of 3 biological replicates per stage are individually plotted. (B) Organ-specific expression in unfed females. (C) Organ-specific expression in engorged females. SG: salivary glands, MG: Midguts, MT: Malpighian tubules. Carcass is remainder of the body after removal of organs (SG, Mg, MT and ovaries). Statistical comparison was done by one-way ANOVA followed by *post hoc* Tukey tests.

We next examined IsAQP1 expression in the major organs of adult females including salivary gland, ovary, midgut, and Malpighian tubules, in both unfed and engorged animals. Among major female organs, IsAQP1 expression was highest in the salivary glands (Fig. 4B, one-way ANOVA with post hoc Tukey test, *P*<0.05). After females fed on blood and became engorged, overall expression fell to 36.5% of the levels of unfed females (Fig. 4B-C) a difference that was not statistically significant with our small sample sizes (one-way ANOVA with post hoc Tukey test, *P*=0.11). Even while expression level dropped, the tissue-specific expression pattern continued to resemble that of unfed ticks (i.e., expression was highest in salivary gland; Fig. 4C, one-way ANOVA with post hoc Tukey test, *P*<0.05).

## Discussion

Here we identify AQP-like genes in *I. scapularis* and use these sequences and others (176 total) to study AQP evolution in arachnids and other arthropods using phylogenetic methods. Consistent with previous work, insect-specific aquaglyceroporins (Entomoglyceroporins, or ‘Eglps’) clustered together within classical water-specific AQPs (Finn *et al*., 2015). In contrast to previous work (Stavang *et al*. 2015), we find that arachnid AQP-like sequences formed a major clade which seems to derive from classical GLPs. More specifically, our analysis placed a majority of arachnid AQP-like sequences (n=68/94), including 51/52 non-Bib AQPs from 13 tick species, into a monophyletic clade nested within GLPs (Figure 2). We also found a few putative classical AQP and even Bib-like genes in arachnids—though none of these has been functionally described. Our findings therefore suggest that in arachnids (and potentially in their closest relatives as well; Ballesteros *et al*., 2022) the GLP gene family has expanded considerably while some (but not all) classical, water-specific AQP genes were lost, perhaps beginning soon after they split from insects and other mandibulates more than 550 million years ago (Giribet and Edgecombe, 2019). Further work—e.g., to carefully inventory and characterize the aquaporins of focal taxa—is needed to understand the evolution of this gene family in arachnids including ticks.

We examined and characterized both the IsAQP1 genomic locus and gene product. We found the IsAQP1 locus to be intron-rich, as expected (Gulia-Nuss *et al*., 2016). Next, we functionally characterized the permeability of IsAQP1 to water, glycerol, and urea. Biochemical characterization showed IsAQP1 has high water channel activity—moving water even more efficiently than a control classical AQP from an insect (Figure 3A). Yet unlike most GLPs, IsAQP1 was a poor conductor of glycerol and was impermeable to urea (Figures 3B-C). Thus, despite being phylogenetically a GLP, IsAQP1 behaves similarly to a classical AQP (though with some remaining glycerol permeability). This finding mirrors a recent study in insects, in which Finn *et al*. described the evolution of GLP-like channels (i.e., permeable to both water and other specific solutes) from classical AQP precursors (2015). Together, our phylogeny and functional analyses imply the converse: that a (largely) water-specific channel—IsAQP1—evolved from a GLP precursor in this arachnid. In other words, arachnids (and especially ticks) may have co-adapted classical GLP homologs to function as water-specific AQPs, an evolutionary path likely related to the depauperate condition of the classical AQP gene family in this clade. Only one other tick AQP, *RsAQP1* from the brown dog tick, *Rhipicephalus sanguineus*, has been biochemically characterized. *RsAQP1* has water channel activity that was inhibited by mercury (Hg^2+^, 100 μM), but it did not have glycerol or urea channel activity (Ball *et al*., 2009). The same author mentioned in a review as unpublished data that another *R. sanguineus* AQP exhibited high glycerol and urea channel activity (Campbell *et al*., 2008). These findings suggest that AQP-like channels may be more evolutionarily labile than was previously assumed, though more studies are needed to functionally characterize AQPs in diverse arachnids to verify the hypothesis that arachnid AQPs are derived primarily from the GLP lineage.

The diversity of AQPs across arthropods may reflect the diverse roles of these channels in maintaining homeostasis and many other aspects of their biology. AQPs function not only in diuresis and desiccation resistance (e.g., Drake *et al*., 2010; Liu *et al*., 2011), but also heat tolerance, intrauterine lactation, and cold tolerance (Benoit *et al*., 2014a; Philip *et al*., 2008, 2011). The unusual osmoregulatory challenges faced by ticks across their life cycle could have shaped the evolution of their aquaporins. Broad scale patterns of evolution may be visible in AQP gene evolution (e.g., Finn *et al*., 2014), but more work is needed to understand the drivers of AQP evolution in arthropods including hematophagous insects and arachnids.

Unlike many AQPs (e.g., Kuwahara *et al*., 1997, Shi and Verkman 1996, Campbell *et al*., 2008), the water channel activity of IsAQP1 was not inhibited by mercury (Figure 3A). Mercury inhibition is thought to rely on the ion binding to specific cysteine residues (Jung et al., 1994; Kuwahara *et al*., 1997; Savage and Stroud, 2007; Hirano *et al*., 2010), though the effects of mercury on AQPs can be complex (e.g., Frick *et al*., 2013). IsAQP1 has three cysteines including Cys^200^, a residue that may be homologous to the verified mercury-sensitive cysteine of other AQPs (e.g., Cys^189^ of human AQP1 and Cys^181^ of human AQP2; Preston et al. 1993), both of which sit 3 residues proximal to the second NPA motif (Figure S1; Shi and Verkman 1996). However, Cys^200^ of IsAQP1 is positioned further upstream from the NPA domain (Figure S1) and this positional shift may explain the loss of mercury sensitivity. Publicly available structural predictions made with AlphaFold2 (Jumper *et al*., 2021, Varadi *et al*., 2021) suggest Cys^200^ may lie within an a-helix while Cys^189^ and Cys^181^ are exposed residues within the channel pore, allowing them to bind mercury. Explicit functional studies would be needed to test this hypothesis—for example, to functionally test the mercury sensitivity of IsAPQ1 with Cys^202^.

Our expression data also shed light on the function of IsAQP1. We detected some IsAQP1 expression at multiple life stages and within multiple tissues, but expression was strikingly enriched in adult female salivary glands (Figure 4B-C). IsAQP1 expression was especially elevated in the salivary glands of un-engorged females (Figure 4C), i.e., expression was highest ahead of an anticipated blood meal. High AQP expression in salivary glands has been noted in other tick species including *Haemaphysalis qinghaiensis* (Niu *et al*. 2022), *Ixodes ricinus* (Campbell *et al*. 2010), and *Rhipicephalus sanguineus* (Ball *et al*. 2009). *IsAQP1* additionally showed low expression in the Malpighian tubules (Figure 4B), but very little expression in gut and ovary—two tissues commonly reported to have high AQP expression in ticks (Niu *et al*. 2022, Ball *et al*. 2009, Holmes *et al*. 2008, Campbell *et al*. 2010). AQPs are typically expressed in a tissue-specific manner (e.g., Tsujimoto *et al*., 2013, Campbell *et al,*. 2008, Van Ekert *et al*. 2016), and expression level may be adjusted dynamically to meet the fluctuating water balance needs of the animal (e.g., Liu *et al*., 2011, Martini *et al*., 2004). Because salivary glands are the major osmoregulatory organ in ticks (Bowman and Sauer, 2004), this expression pattern suggests that IsAQP1 functions in water balance during blood feeding. Indeed, while feeding on hosts, ticks must return 70% of the water and ions from acquired blood back into the host (Bowman and Sauer, 2004), which implies a need for an efficient water channel such as IsAQP1.

We found high variability between replicates when examining IsAQP1 expression across life stages in ticks sourced from BEI Resources. The source of this expression variation is unclear, but it could be related uncontrolled variation in age, housing, or shipping conditions. Repeating this work using animals kept in controlled lab conditions would be necessary to determine if this variation arises from inherent biological sources or from environmental perturbation.

Currently, tick control primarily involves the use of chemical acaricides, yet these tools lead to the evolution of resistance and environmental contamination (e.g., Murgia *et al*., 2019), leaving us in urgent need of new methods of tick control. Our work suggests tick aquaporins, including IsAQP1, may be effective molecular targets for new control efforts aimed at disrupting essential biological processes in ticks. Because pathogens are passed from ticks to hosts via saliva, the aquaporins involved in salivation remain attractive targets for the development of novel control methods. Indeed, aquaporins continue to gain attention as targets of anti-tick vaccination strategies (e.g., de la Fuenta *et al*., 2016, Guerrero *et al*., 2017, Scoles *et al*., 2022). We attempted to knock down IsAQP1 expression using RNAi for *in vivo* functional characterization that could give insight into potential control targets, but gene knock-down was unsuccessful (data not shown).Thanks to the recent development of new methods for targeted CRISPR genetic manipulations this species (Sharma *et al*., 2022), future research could functionally test the importance of this gene in tick physiology and explore its potential for use as a tool of vector control.

## Experimental Procedures

All methods were carried out in accordance with Penn State University institutional guidelines and regulations.

### Animal and tissue handling

Ticks were obtained from both the field and from BEI Resources. Adult *Ixodes scapularis* were collected in State College, PA using a standard canvas-dragging method. Eggs, larvae, nymphs, male and female adults and engorged female *Ixodes scapularis* were obtained from BEI Resources (NIAID, NIH). Field-collected ticks were used for cloning and sequencing IsAQP1, while purchased ticks were used for tissue- and life stage-specific expression analyses.

For cloning and sequencing, field-collected ticks were either processed immediately or stored at –80 °C until processing. For gene expression analysis, (including organ dissection) ticks were kept alive at room temperature (22-25 °C) and ambient conditions for 0-4 days prior to processing. Organs (salivary glands, midgut, Malpighian tubules, and ovaries) were dissected in PBS, pooled, and either immediately processed or stored in TRI Reagent (Sigma Aldrich, Saint Louis, MO) at –80 °C until use. Pools were composed as follows: 120-160 individual larvae, 25 nymphs, 8-20 adult males (whole animals or organs), and 8-10 adult females (whole animals or organs). A volumetrically similar clump of eggs comprised a replicate of pooled eggs.

### RNA extraction, cDNA synthesis, and RACE

Whole ticks or dissected organs were homogenized in TRI Reagent (Sigma Aldrich, Saint Louis, MO) in a 1.5 mL microcentrifuge tube with a pestle (Kimble Kontes, Fisher Scientific, Hampton, NH). Whole RNA was extracted following the manufacturer’s instructions. Extracted RNA was treated with DNase (TURBO DNAfree, Life Technologies, Carlsbad, CA) for 1 hr at 37 °C. First-strand cDNA synthesis was performed using Accuscript High Fidelity Reverse Transcriptase (Agilent Technologies, Santa Clara, CA) with oligo d(T)_20_ primer on 1 μg of total RNA at 42 °C for 2 hr.

We used RLM-RACE (**R**NA **L**igase-**M**ediated **R**apid **A**mplification of **c**DNA **E**nds) to determine the unknown sequence at the 5’ and 3’ ends of IsAQP1 cDNA as previously described (Tsujimoto *et al*., 2013). Briefly, RNA was extracted from adult ticks (both male and female adults) using TRI Reagent. RNA processing for RACE was done according to the manufacturer’s instructions (GeneRacer, Life Technologies, Carlsbad, CA). RACE PCR was performed with gene-specific primers based on the predicted gene sequence at VectorBase.org (ISCW003957) plus an anchor primer (3’ RACE) or GeneRacer 5’ primer/GeneRacer 5’ nested primer (5’ RACE) (Table S1). RACE PCR products were separated by agarose gel electrophoresis, and bands were excised for gel extraction using a Gel Extraction Kit (QIAGEN, Valencia, CA). Gel-extracted PCR products were cloned into pJET1.2/blunt vector using CloneJET PCR Cloning Kit (Fermentas, Glen Burnie, MD). The inserts were then Sanger sequenced, and the resulting full length cDNA sequence was submitted to GenBank (accession number KT988052).

### Mapping exons in genomic sequence

We mapped the IsAQP1 cDNA sequence to the genome to determine the location and structure of the gene locus. We completed a BLASTn search of the *I. scapularis* genome sequence at VectorBase (vectorbase.org) using the full-length cDNA as query. We then manually adjusted exon-intron boundaries so that all introns have a canonical splice signal (GT-AG) and submitted our revised annotation to VectorBase. Visualization of the IsAQP1 locus was accomplished using the web-based software FancyGene (Rambaldi and Ciccarelli, 2009).

### Phylogeny estimation

We obtained AQP sequences from public sources (e.g., GeneBank, FlyBase, VectorBase, transcriptome shotgun assemblies, whole genome shotgun contigs). Sequences were identified via arthropod-specific searches for ‘aquaporin,’ BLAST searches using arthropod aquaporin sequences as query, and from previous studies (Stavang *et al*. 2015, Niu *et al*. 2022). Scorpion transcriptome contigs were provided by Dr. Thorsten Burmester (Zoological Institute and Museum, University of Hamburg, Germany; Roeding *et al*., 2009). All taxon and accession information is given in Table S1. Polypeptide sequences (translated from nucleotide sequence as needed at expasy.org) were used as input, and all sequences ≥200 amino acids with at least one NPx motif (where x = any amino acid) were included in phylogeny estimation. Sequences were aligned using SeaView (Gouy *et al*., 2010) built in Clustal Omega, where unaligned regions were filtered using the “Gblocks” option with the least stringent options. The phylogenetic tree was constructed using maximum likelihood in PhyML 3.1 with 500 bootstraps, and plotted using FigTree v1.4.4.

### Real-time quantitative PCR (qRT-PCR)

We used qRT-PCR to quantify the expression of IsAQP1 in different life stages and tissues. Primers were designed using Primer3 (Koressaar and Remm 2007; Untergasser *et al*., 2012), and designed to amplify 53-145 bp fragments of the cDNA (Table S2). We empirically verified that all primer pairs have an amplification efficiency (E) between 0.9 and 1.1 using the same qPCR protocol. cDNA was diluted to 1/50 in nuclease-free H_2_O, of which 2.5 μL was used for 10 μL reactions using the RotorGene Q qPCR system (QIAGEN). Reactions were performed with 5 min at 95 °C, followed by 45 cycles of 95 °C for 5 sec, 60 °C for 10 sec, and melt curve analysis from 50 °C to 95 °C. Expression was calculated relative to a housekeeping gene (Glyceraldehyde 3-phosphate dehydrogenase, GAPDH) by the Pfaffl method (2001).

### Cloning, expression in *Xenopus laevis* oocytes, and osmotic swelling assays

We used *X. laevis* oocyte swelling assays to evaluate the channel properties of the cloned *I. scapularis* AQP-like gene. The sequence was cloned as previously described (Tsujimoto *et al*. 2013). Briefly, the complete coding sequence was then amplified using primers with a restriction site at the 5’ end (MfeI for upstream primers and NheI for downstream primers). The resulting PCR products were initially cloned into pJET1.2/blunt vector (Fermentas, Glen Burnie, MD) and after restriction digestion of the cloned insert, the *IsAQP1* CDS was ligated into EcoRI/NheI-treated pXβG-myc. The purified construct was linearized using XbaI for *in vitro* complementary RNA (cRNA) transcription with T3 RNA polymerase (Agilent Technologies, Inc., Santa Clara, CA). Synthesized cRNA was purified with the RNeasy Mini kit (Qiagen), and its integrity was analyzed by denaturing agarose gel electrophoresis.

For comparison controls, we additionally evaluated AQP genes AgAQP1B, ClAQP1 and ClAQP1/ClGlp1 co-expression (Tsujimoto *et al*., 2013, 2017) using the same methods, allowing for direct comparison of channel properties. *Xenopus laevis* ovaries were purchased from Xenopus I (Dexter, MI), defolliculated with collagenase I (Sigma) and injected with 50 nL of 100 ng/μL cRNA solution (5 ng) or 50 nL of nuclease-free water as control. cRNA-injected oocytes were incubated for three days in ~200 mOsm Modified Barth’s Solution (MBS: 88 mM NaCl, 1.0 mM KCl, 2.4 mM NaHCO_3_, 15 mM Tris, 0.8 mM MgSO_4_, 0.4 mM CaCl_2_, 0.3 mM Ca(NO_3_)_2_, pH 7.6, containing 0.5 mM theophylline, 100 U/mL penicillin and 100 μg/mL streptomycin) at 16 °C to allow for overexpression of aquaporin in the oocyte plasma membrane. The oocytes were then subjected to a swelling assay as previously described (Tsujimoto *et al*., 2017). Briefly, we transferred oocytes to MBS diluted to 70 mOsm. The change of oocyte volume was monitored at room temperature using a digital camera (Olympus DP-21) equipped stereo-microscope (Olympus SZX-16) for 60 seconds with the relative volume (*V/V*_0_) calculated from the cross-sectional area at the initial time (*A*_0_) and after the specified time interval (*A_t_*): *V/V*_0_ = *(A_t_/A_0_)^3/2^*. The coefficient of osmotic water permeability (*P_f_*) was determined from the initial slope of the time course [*d(V/V_0_)/dt*], average initial oocyte volume (*V*_0_ = 9 × 10^-4^ cm^3^), average initial oocyte surface area (*S* = 0.045 cm^2^), the molar volume of water (*V_w_* = 18 cm^3^/mol), and the osmotic solute gradient (osm_in_ - osm_out_: *P_f_* = (*V*_0_ × *d(V/V*_0_)/*dt*)/(*S* × *V_w_* × (osm_in_ - osm_out_)). A minimum of six (range 6-12) individual oocytes were tested for each group.

To test the effect of mercury (Hg^2+^) on water permeability (Preston et al. 1992), oocytes were placed in 500 μM HgCl_2_ in MBS for 5 min prior to the swelling assay. We tested the reversibility of Hg^2+^ treatment by transferring Hg^2+^-treated oocytes (500 μM, 5 min) to 5 mM 2-mercaptoethanol in MBS for 10 min.

Glycerol and urea permeability were also assessed in separate assays by replacing diluted MBS (i.e., 70 mOsm) with MBS that was mixed with an equal volume of iso-osmotic (200 mOsm) glycerol or urea solution. Data were compared by one-way ANOVA followed by Tukey’s correction for multiple comparisons. All analyses were completed using Graphpad Prism software.

## Acknowledgements

This study was supported by USDA Hatch funds (Project PEN04769), a grant with the Pennsylvania Department of Health using Tobacco Settlement Funds, and funds from the Dorothy Foehr Huck and J. Lloyd Huck endowment to JLR, NSF/BIO grant 1645331 to JLR and JMS, and NIH/NIAID grant R01AI080615 to UP. BEI Resources (NIAID, NIH) provided *Ixodes scapularis* used in this research (NR-42510). We thank Dr. Thorsten Burmester (Zoological Institute and Museum, University of Hamburg, Germany) for providing the contig set from the scorpion transcriptome.

## Data Availability Statement

IsAQP1 mRNA sequence has been deposited in GenBank with an accession number K988502.

## Abbreviations

AQP: aquaporin
C: carcass (rest of the body after SG, Mg, MT and Ov removal)
CDS: coding sequence
cRNA: complementary RNA
GLP: aquaglyceroporin
IsAQP1: *I. scapularis* aquaporin 1
Mg: Midguts
MT: Malpighian tubules
mOsm: milliosmolar
NPA: asparagine-proline-alanine
Ov: ovaries
Pf: osmotic water permeability
Ps: solute permeability
qRT-PCR: quantitative real-time PCR
SG: salivary glands
UTR: untranslated region

## Tables

**Table S1:**
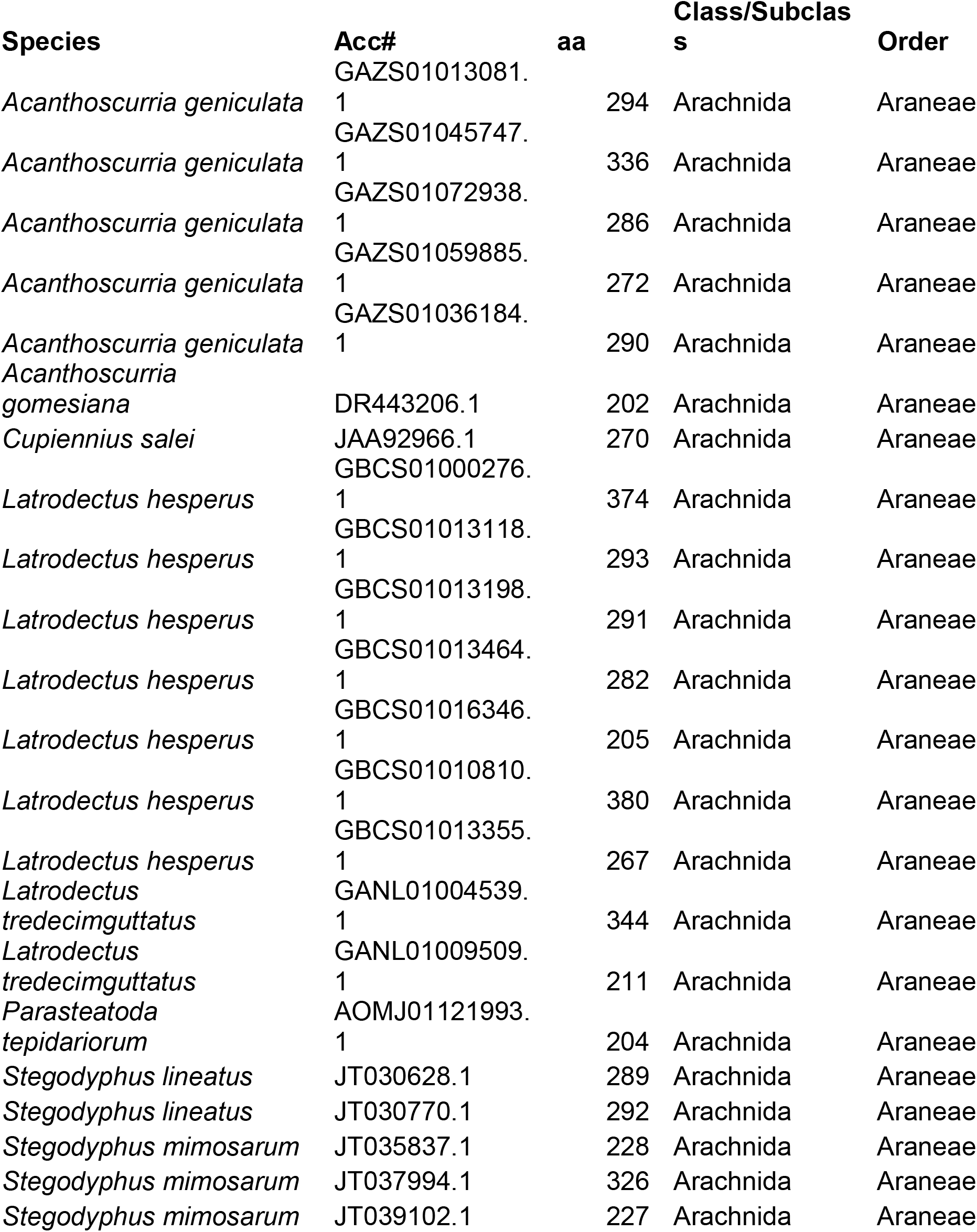
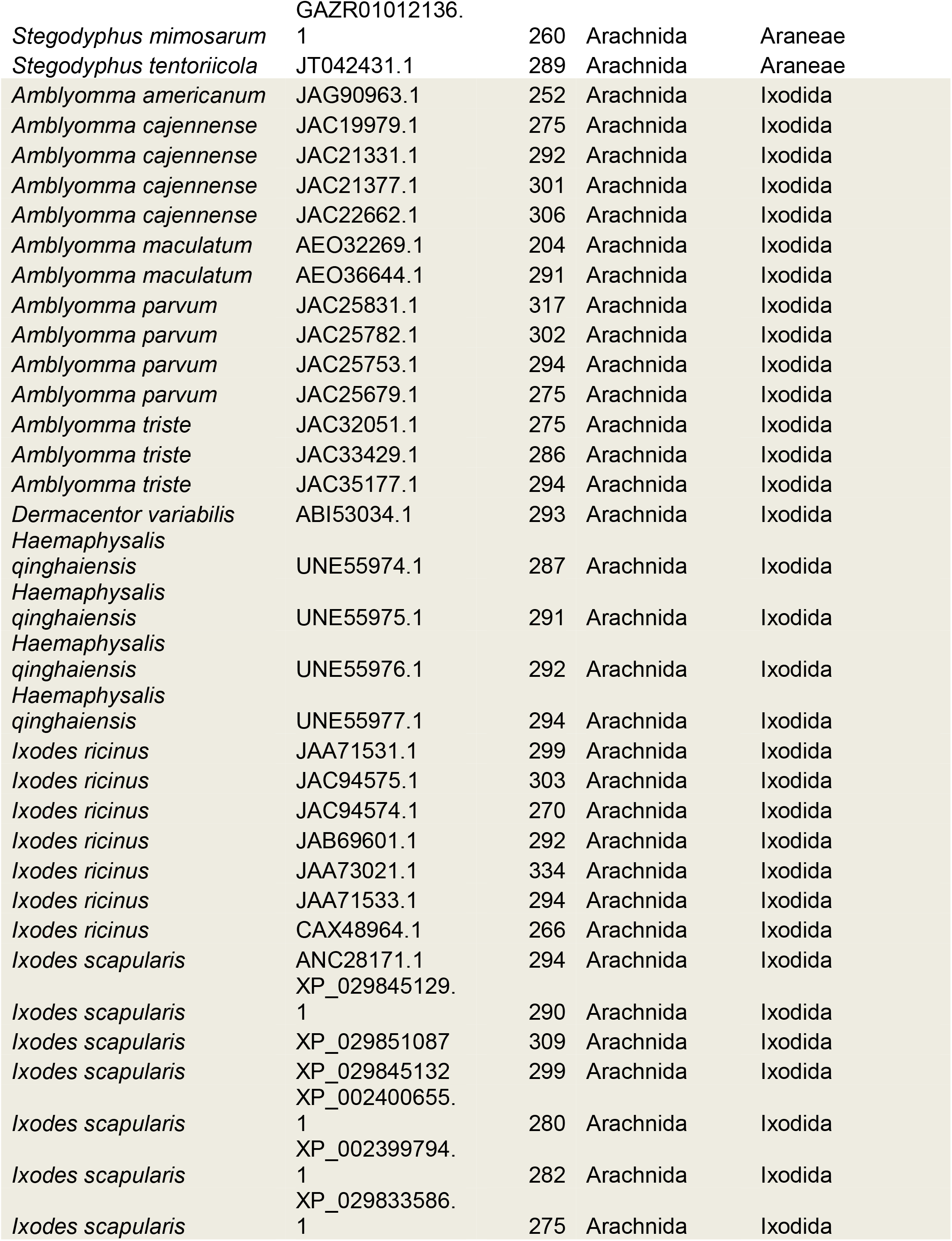
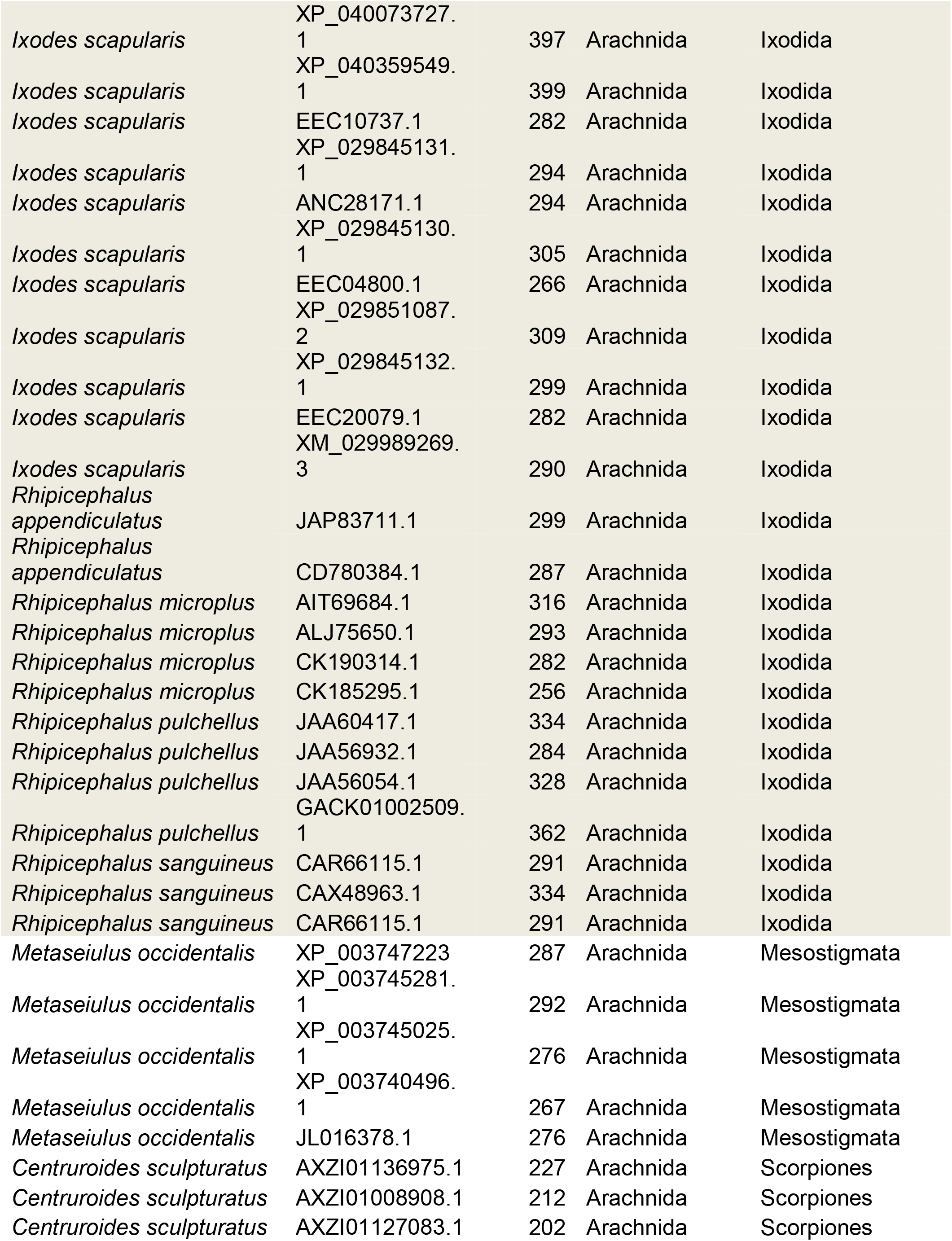
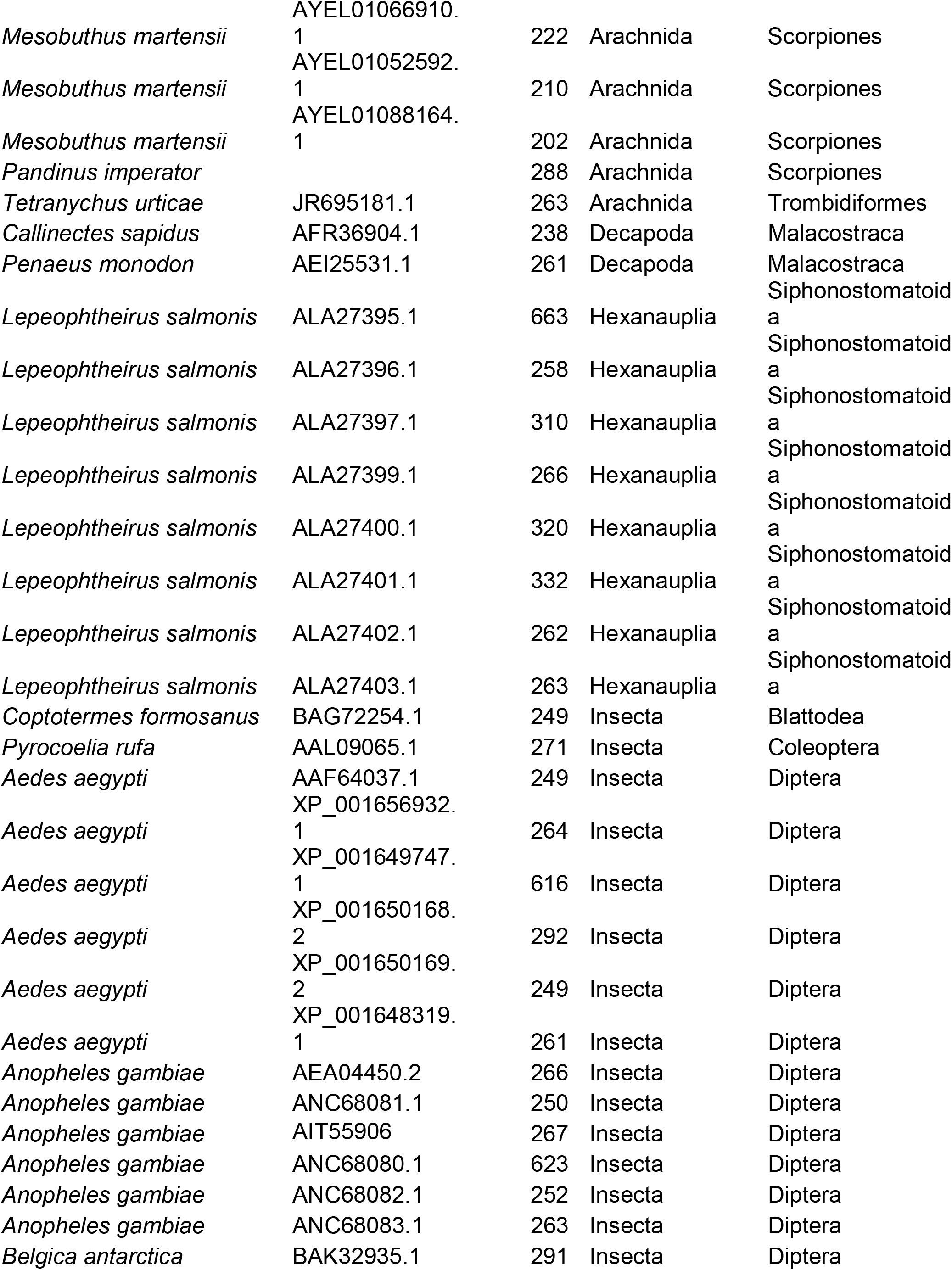
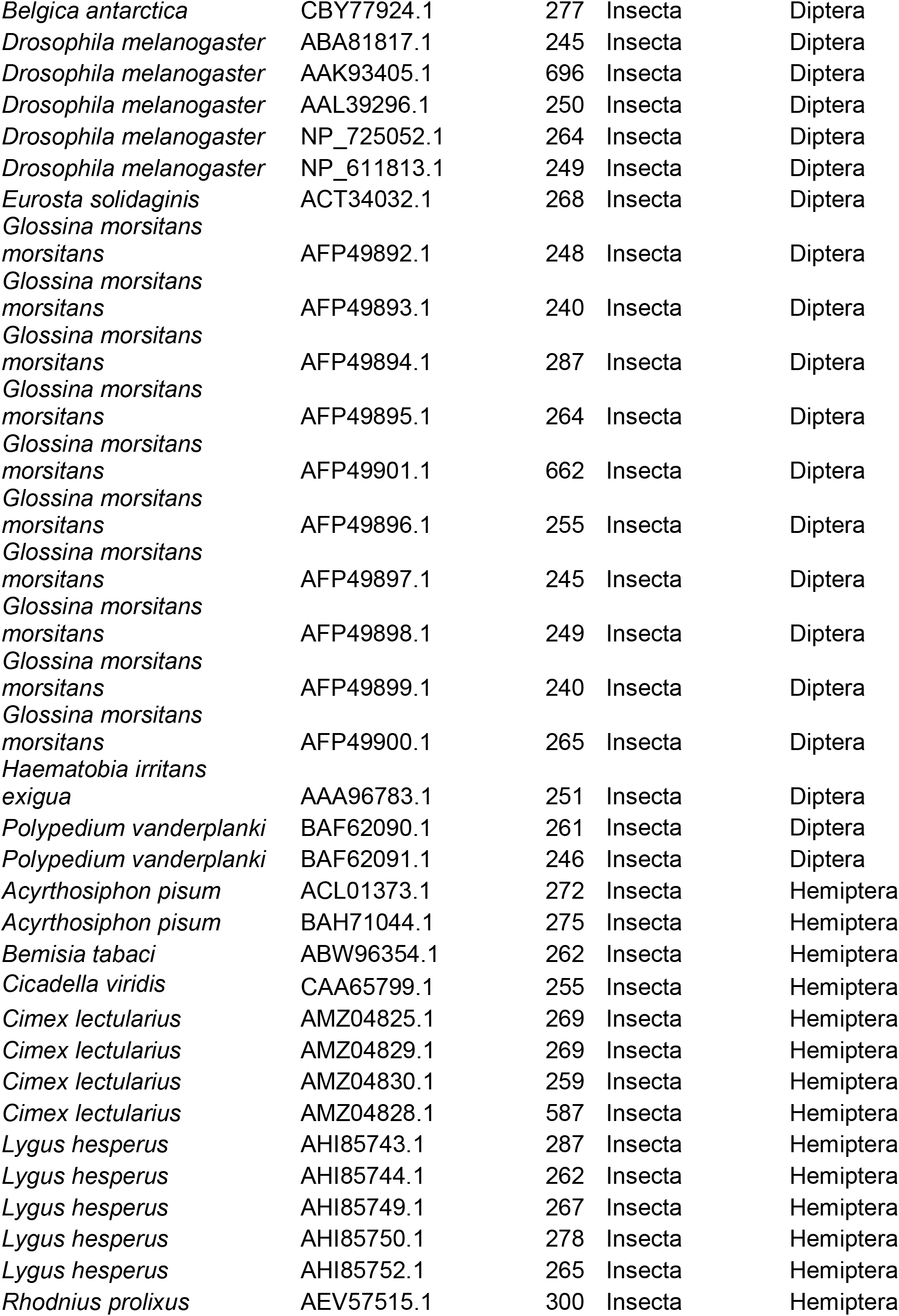
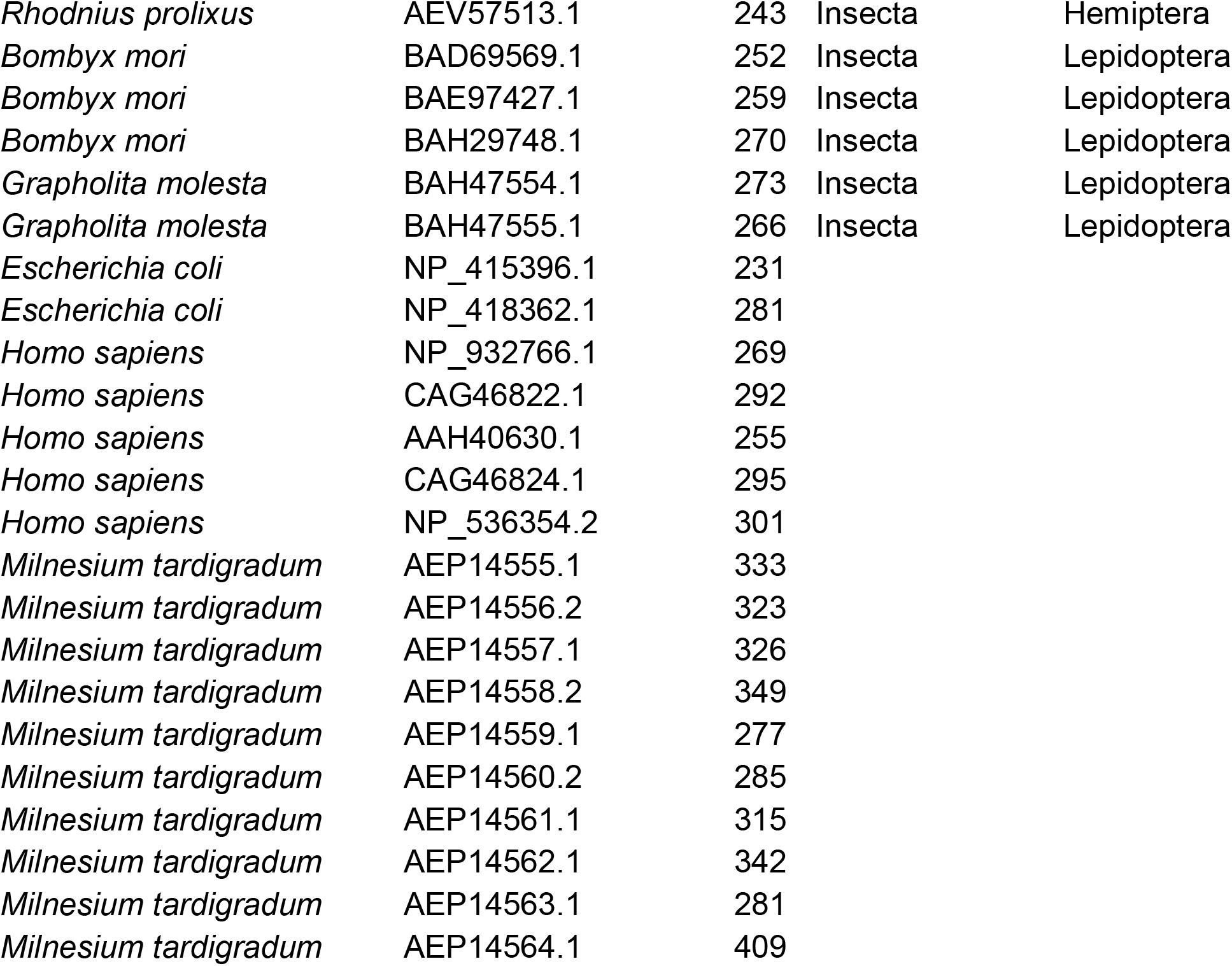
Names of AQP-like genes, species of origin and GenBank accession numbers for the amino-acid sequences used in Fig. 2.

**Table S2:**
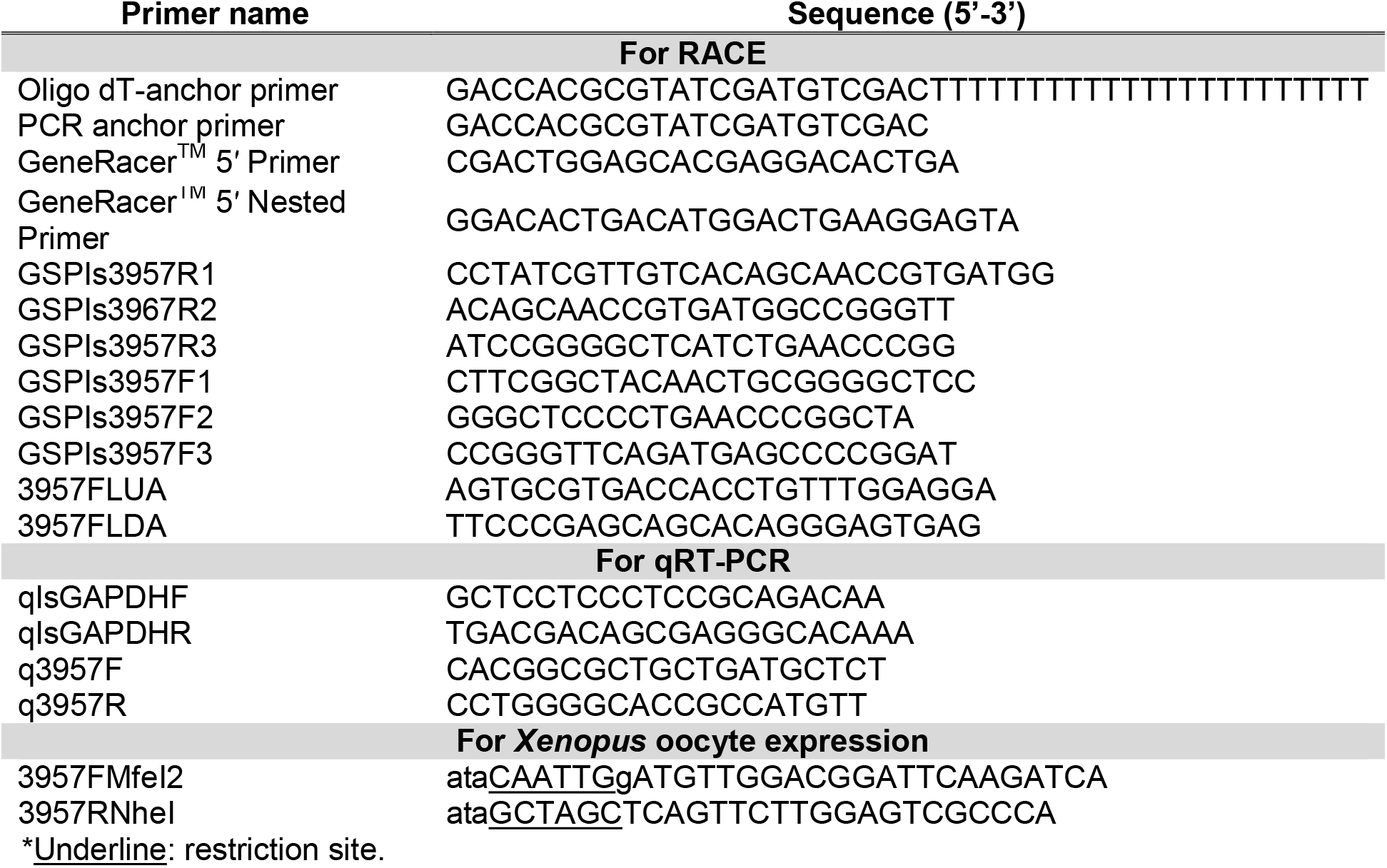
primers used for this study.

## Supplemental Figure

**Figure S1.**
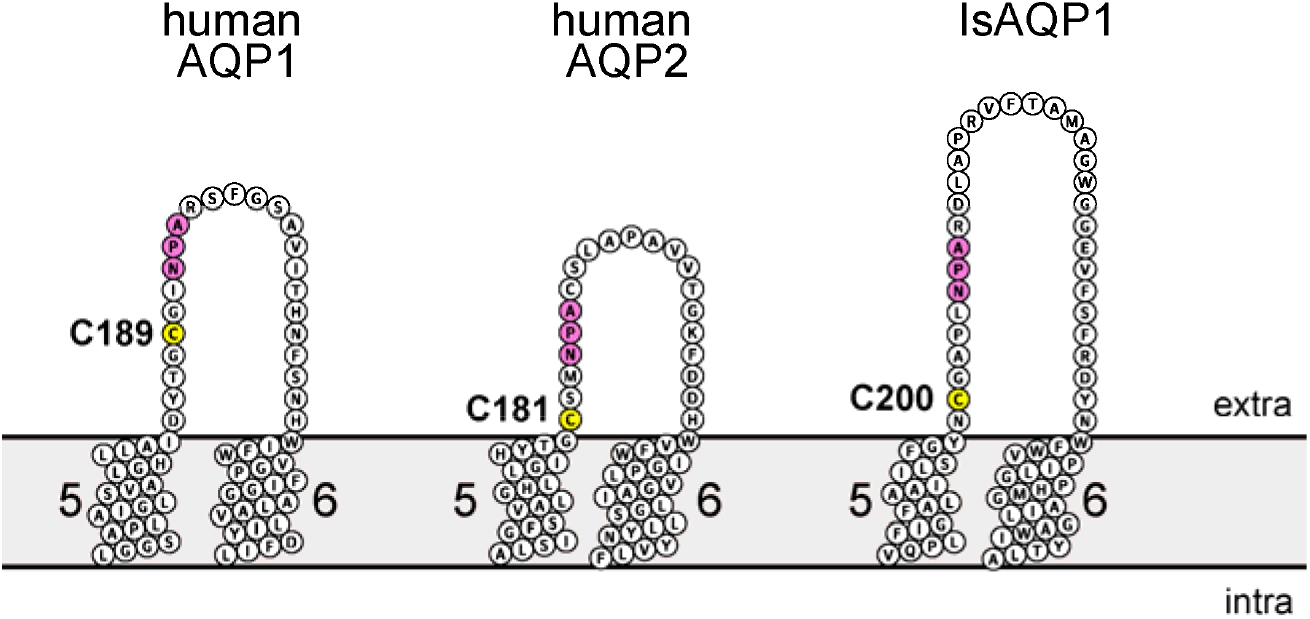
Comparison of cysteine residue (yellow) position relative to the second NPA domain (pink) in the mercury-sensitive AQPs human AQP1 and AQP2, and the mercury-insensitive AQP IsAQP1 (right). Truncated Protter diagrams are shown. The 5^th^ and 6^th^ transmembrane domains of each polypeptide are labeled, as are the extra- and intra-cellular sides of the membrane.

